# T6SS-mediated competitive exclusion among *Pantoea agglomerans* associated with plants

**DOI:** 10.1101/2023.03.17.533219

**Authors:** Kai Ripcke, Devani Romero Picazo, Shreya Vichare, Daniel Unterweger, Tal Dagan, Nils F. Hülter

**Affiliations:** Institute for General Microbiology, Kiel University, 24118, Kiel, Germany; Institute for Experimental Medicine, Kiel University, Michaelisstraße 5, 24105 Kiel, Germany; Max Planck Institute for Evolutionary Biology, August-Thienemann-Straße 2, 24306 Plön, Germany

**Author notes:** To whom correspondence should be addressed: Nils F. Hülter, Kiel University.

## Abstract

Establishment of symbiotic interactions with a host habitat depends on colonization success of the microbiome members. One route to success is increasing competitiveness against genotypes having similar adaptations. Many bacteria deploy the type VI secretion system (T6SS) to deliver toxic effector proteins into the cytoplasm of competing cells, whereas the attacking cells are protected from self-intoxication by immunity proteins. While the evolution of antagonistic interactions between species competing for similar niches is expected, the interactions between closely related strains having a similar core genetic makeup, remains understudied. Here we show that the strength of T6SS-mediated competition between closely related *P. agglomerans* strains is not associated with phylogenetic relatedness and rather depends on their effector and immunity genes repertoire. Annotating the T6SS components in eleven *P. agglomerans* isolates from wheat seeds, we observed a heterogeneous composition of effector and immunity determinants. Competition experiments further showed profound differences in the strain survival following reciprocal T6SS-mediated interactions. Reconstructing the evolution of T6SS loci across plant-associated *P. agglomerans* isolates indicates intra-chromosomal recombination as the main driver of effector and immunity gene diversification. Our results provide empirical data on intraspecific interactions during microbiota assembly likely to be at play during colonization of germinating plant seeds.

## Introduction

The ecological principles that govern microbial community assembly processes depend on inter- and intra-species interactions, even in host-associated habitats where the host organism actively selects the composition of its microbes. Host-associated habitats offer different niches in which resources, whether nutrients or unoccupied space, are finite. Hence for most bacteria in symbiosis, the pursuit of reproductive success is not just a function of metabolic efficiency, versatility, or niche specialization in the densely populated, species-rich host environment. Bacteria have evolved a multitude of strategies and mechanisms to proliferate under competitive circumstances with other bacteria. Bacteria deploy a wide range of enzymes and transporters to gain access to nutrients in an exploitative fashion to gain reproductive benefits. Competitiveness and thus evolutionary success, can be achieved through cooperative or synergistic resource utilization. An alternative for synergistic resource utilization is to inhibit the competitor proliferation and thus avoid competition. Indeed, bacteria have evolved diverse warfare mechanisms; this includes the utilization of contact-independent and contact-dependent mechanisms for harming and killing competitor cells. The ubiquity of mechanisms for interference competition (or bacterial warfare) in bacterial communities is evidence of their importance as drivers of ecological and evolutionary success [1].

The arsenal of bacterial weapons is plentiful and bacterial organisms typically harbor multiple systems [2,3]. Contact-independent mechanisms include e.g., chemical warfare, where bacteria utilize secretion of diffusible compounds such as toxins and antimicrobial peptides via secretory pathways and pores that target predominantly an adversary’s ability to grow and divide [4]. Among the systems of contact-dependent combat strategies is the widely distributed type VI secretion system (T6SS) [5–7]. The system constitutes a toxin-delivery apparatus with a toxin-equipped syringe-like cell wall puncturing needle that is expelled from a spring-loaded contractile sheath upon induction. The T6SS is, in principle, a combination of a toxin and immunity mechanism. In addition to the genes encoding the structural components, T6SS systems encode both toxic effectors and cognate immunity proteins [8], while effectors are translocated into the cytoplasm of the attacked cell, the immunity proteins remain in the attacking cell where they prevent self-intoxication and provide immunity to attacks from isogenic cells or cells deploying an identical effector. The effectiveness of T6SS systems generally depends on the repertoire of effector and immunity genes that are present in a particular system. Cells lacking immunity against the effectors translocated into their cytoplasm during attack can be killed.

Microbial diversity in a given environment is usually assessed via the number of distinguishable species and their abundance. Nonetheless, it has become evident in past years that microbial communities comprise multi-strain assemblages of the resident species. For example, in the deep-sea mussel environment, multiple sulfur-oxidizing symbiont strains coexist within individual hosts [9]. Additionally, closely related *Vibrionaceae* strains are shown to differentially contribute to the polysaccharide breakdown process in the marine environment [10]. Genotypes in such multi-strain assemblages generally exhibit high degrees of similarity across their core genetic make-up (i.e., within the limits of accepted variability on the species level), but can display striking differences across their accessory genome and hence specific phenotypes [11]. Considering the mechanistic basis of T6SS, spiteful interactions between closely related strains sharing immunity profiles are expected to be even. Conversely, but intuitively, inclusive fitness theory argues that the evolution of spitefulness between related strains should prevail, when these have overlapping metabolic requirements and share an ecological niche. Studies on the ecological implications of competitive interactions between organisms having similar killing mechanisms – especially on the intraspecific level – for microbial community assembly in nature yielded contrasting conclusions. For example, bacteriocin-mediated interactions between populations of related *Pseudomonas fluorescens* species from soil indicated no relationship between the level of inhibition and the genetic similarity between competitors [12]. In contrast, most likely shared resistance mechanisms prevent among-strain inhibition in populations of the entomopathogenic bacterium *Xenorhabdus bovienii* [13], while spiteful behavior between *Pseudomonas aeruginosa* isolates appeared predominantly between genotypes of “intermediate” genetic distance [14]. Speare et al. (2018) [15] and Kostiuk et al. (2017) [16], along with [17–19], have provided evidence for intraspecific competition mediated by the T6SS in *Vibrio cholerae*. In this organism, the presence or absence of effector/immunity gene pairs determines the compatibility of different strains. Immunity proteins are effective only against effectors of the same kin type, while strains carrying different effector genes are not protected. Consequently, genetic differences in effector/immunity gene pairs within an otherwise isogenic T6SS module play a decisive role in the behavior of strains that otherwise exhibit similar fitness in their habitat. However, it remains unclear whether related strains exhibit similar competitive exclusion interactions, or if such interactions are associated with their evolutionary relatedness.

Here we study the strength of competitive exclusion among closely related *Pantoea agglomerans* strains and evaluate the level of pairwise inhibition in the context of strain divergence. Strains belonging to the *P. agglomerans* species are commonmembers of the epiphytic and endophytic microbial assemblages of plants [20,21]. While some strains exhibit pathogenic potential, many species members are considered to be plant growth-promoting bacteria [22–26]. In a previous study, we have isolated eleven *P. agglomerans* strains from seeds of wheat cultivars and landraces [27]. Using an *in vitro* bottom-up approach we assessed the strength of T6SS-mediated competition among these strains. Furthermore, we quantified the genetic divergence of the tested strains and compared it with the outcome of cross-strain growth inhibition. Our study supplies novel insights on the evolution of antagonistic interactions among closely related strains in plant host-associated microbial communities.

## Results

### The repertoire of T6SS effector and immunity genes in *Pantoea agglomerans* strains

Components of the T6SS have been previously described in several *Pantoea* species including *P. agglomerans* [28–30] and *Pantoea ananatis* [31]. Using the known genes as queries we identified gene clusters that putatively encode T6SS components and examined their conservation among eleven *P. agglomerans* strains isolated from seeds of wheat cultivars (strains A2, B1, B3, R1, and R5) and landraces (strains T1 to T6). All of the eleven strains included up to three distinct T6SS gene clusters (Fig. 1A), in line with previous reports of the T6SS in the genus *Pantoea* [28].

**Figure 1.**
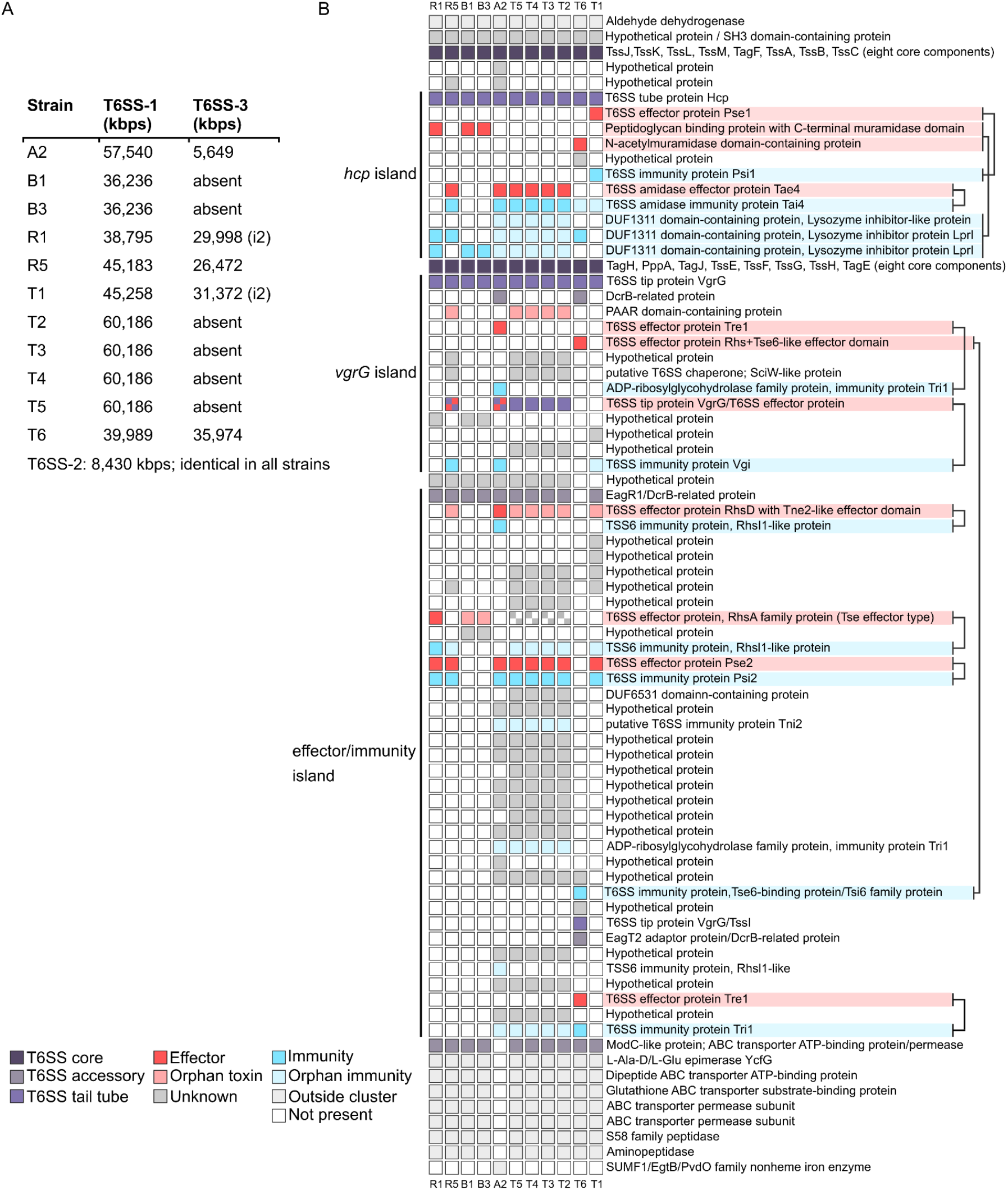
T6SS clusters in P. agglomerans and a detailed view of cluster 1. **(A)** T6SS cluster 1 and 3 identified in the eleven *P. agglomerans* strains. All clusters, except cluster 3 in strain R1 and T1, belong to the subtype i3 of T6S systems [37,38]. Detailed information on gene content and structure of the three clusters is provided together with the respective locus tags in Supplementary Table S1. **(B)** Genetic organization of T6SS cluster 1 in the eleven *Pantoea agglomerans* isolates from wheat seeds. The location of the genes within the cluster is shown in the form of a gene presence/absence pattern. Predicted effector/immunity pairs are indicated.

Here we number the T6SS clusters according to their gene content similarity to previous studies [28,29]. Most of the genes encoding the core T6SS components were conserved throughout the sampled set across three different genomic regions. Examples are, the baseplate subunit TssM, the tube protein Hcp, and the ATPase TssH, which are encoded in cluster 1 (Fig. 1B and Supplementary Table S1). The pattern of gene presence and absence in cluster 1 furthermore revealed two variable regions that are typical for genomic loci encoding for effector and immunity gene pairs as previously described for *P. agglomerans*.

Across all eleven strains, cluster 1 displays the architectural features observed by Carobbi et al. (2022) [29] and Boyer et al. (2009) [32] and can be divided into five major subset regions of genes. Cluster 1 starts with a SH3-domain-containing protein (similar to A7P61_RS17235 in *P. agglomerans* pv. *betae* strain 4188) and continues into the conserved region that comprises the protein encoding the core T6SS genes *tssJ* to *tssC*. This conserved part is followed by the *hcp* island with flexible gene content and the second conserved region, which spans the genes encoding *tagH* to *tagE*. This conserved region is followed by the *vgrG* island and a very flexible region, which we term effector/immunity island. This region presumably codes for several effector and immunity gene pairs, not all of which we could predict a function. Our analysis of the T6SS cluster 1 led us to conclude that the cluster extends even beyond the effector/immunity gene pair *pse2*/*psi2*, which was set at the 3’ end of the T6SS cluster 1 in *P. agglomerans* pv. *betae* [29]. We identified downstream of the *pse2*/*psi2* effector/immunity gene pair several genes that we predict to be part of the cluster. The gene we consider to be the first gene outside of cluster 1 is *ycjG*, which encodes a L-Ala-D/L-Glu epimerase, while in strain A2 the first gene outside the cluster 1, for which we could not predict a T6SS-associated function, is a gene encoding a SUMF1/EgtB/PvdO family nonheme iron enzyme (Fig. 1B).

The auxiliary cluster 2 has an identical size and gene composition in the eleven isolates (see Supplementary Table S1). Cluster 2 comprises homologs of both core and accessory T6SS genes found in cluster 1, including *tagE, tagH, tagF*, and *tssM*. Components of both clusters were suggested to facilitate the assembly of a derivative secretion apparatus - its existence and function remain to be shown [31].

Cluster 3 exhibits greater variability in both gene content and architectural properties (see Supplementary Table S1 and Supplementary Figure S4). This cluster is absent in strains T2 to T5, while strains B1 and B3 retain a 5’-terminally truncated copy of a presumably T6SS-related gene, *vasL* (ImpA family of inner membrane proteins), at the respective genomic location. Despite this variability, conserved genes within Cluster 3 largely exhibit a matched synteny. These genes include additional homologs of the twelve structural component genes (*tssA*-*tssG, tssI*-*tssM*), although their presence across strains is heterogeneous. Strains R1, R5, and T1 have a complete set of the core genes, suggesting that they may possess a functional arsenal for establishing an alternatively regulated T6SS. Cluster 3 in strain T6 exhibits notable differences in location and architecture compared to the other strains. Firstly, it is situated in a distinct genomic context (see Supplementary Table S1 for details). Additionally, the synteny of its core components varies significantly from those in other strains. Despite these dissimilarities, the cluster still encodes the essential core components required for an alternative T6SS system. As noted by Shyntum et al. (2014) [31], the presence of cluster 3 in *P. ananatis* appeared limited to isolates obtained from symptomatic plant material. In the case of *P. agglomerans*, it is not generally considered pathogenic to plants, which raises the possibility that cluster 3 may be more indicative of host specificity rather than pathogenicity.

To identify genes encoding effector and immunity proteins, we first compared our strains with the effector/immunity gene repertoire of *P. agglomerans* pv. *betae* [29], which includes a set of two newly experimentally validated effector/immunity protein pairs: (i) a specialized VgrG effector protein having a C-terminal glucosaminidase toxin domain along with its cognate immunity protein Vgi, and (ii) the effector protein Pse2, which targets the periplasmatic space but whose exact mode of action is not yet understood, together with its immunity protein Psi2. Next, we annotated genes in the variable region of cluster 1 using protein sequence and domain similarity to known effector and immunity genes. Our analysis uncovered nine pairs of effector and immunity genes and several putative orphan-like effectors and immunity genes (Fig. 1B and Supplementary Table S1). In the variable region between the T6SS tube protein encoding gene *hcp* and the gene *tagH*, we confirmed the presence of the T6SS effector/immunity gene pair *pse1*/*psi1* similar to *P. agglomerans* pv. *betae*. This gene pair is only present in strain T1 and absent in the other ten strains. The effector protein Pse1 has strong similarity to pore-forming colicin toxins, while its cognate immunity protein Psi1 is characterized by a transmembrane helix domain. The *pse1*/*psi1* gene pair is followed by the effector/immunity gene pair *tae4*/*tai4* in six of our strains (i.e., A2, B1, T2, T3, T4, and T5). The Tae4 effector protein is a periplasmic active cell wall amidase, which is inactivated by a Tai4 homodimer, resulting in a heterotetrameric inhibition complex in which one Tai4 homodimer can inhibit two Tae4 enzyme molecules [33,34].

Using translated nucleotide queries as input, we queried HHpred [35] and InterProScan [36], to identify the presence of other effector/immunity gene pairs within cluster 1. Within the *hcp* island, we identified two putative peptidoglycan-targeting effector encoding genes. The first gene is present in strains R1, B1, and B3 (e.g., GQQ16_RS01780 in strain R1). We named this effector Pme (*<u>P</u>antoea* muramidase <u>e</u>ffector). The protein displays strong similarity to bacterial transpeptidases on its N-terminus, while the C-terminal region contains a muramidase domain (98.25% probability of similarity between Pme amino acids 130–273 and amino acids 197–334 of the pesticin toxin (PDB: 4AQN) from *Yersinia pestis*). The second effector is only present in strain T6 (GQQ23_RS09490) and encodes a N-acetylmuramidase domain-containing protein. We named this putative effector Ame for (<u>A</u>cetyl<u>m</u>uramidase <u>e</u>ffector). Ame shares similarity with a phage endolysin (100% probability of similarity between Ame amino acids 14–277 and amino acids 29–284 of the endolysin of phage *Escherichia coli* phage FTBEc1 (PDB: 7RUM)). These two effector coding genes likely complement the three sparsely distributed genes that each encode the lysozyme inhibitor domain LprI as already observed in the *hcp* islands of other *Pantoea* strains [29].

Within the *vgrG* island (Fig. 1B), we located the specialized VgrG/Vgi effector/immunity pair described previously [29] in only two of our strains (i.e., A2 and R5). In the strains T5, T4, T3 and T2 the VgrG homologs contain an unknown C-terminal region compared to the specialized VgrG effector in strain A2 and R5, which harbors a domain with similarity to a N-acetylmuramoyl-L-alanine amidase (InterPro: IPR002901) with 98.5% probability of similarity between amino acids 671–829 and amino acids 49–209 of protein 5T1Q (PDB). We note that strains T5, T4, T3 and T2 harbor no *vgi* gene. Upstream of the *vgrG* we located in strain A2 the Tre1/Tri1 effector/immunity pair encoding genes (i.e., GQQ13_RS03745 and GQQ13_RS03740). The Tre1/Tri effector/immunity protein pair shares high identity with the NAD(+)--protein-arginine ADP-ribosyltransferase Tre1 and the ADP-ribosylarginine hydrolase Tri1 (62.6% and 63.2% identity (99% coverage), respectively) from *Serratia proteamaculans* (UniProtKB: A8GG78.1 and A8GG79.1). In the same position, strain T6 (which lacks the *vgrG*/*vgi* gene pair) encodes a protein (GQQ23_RS09565) containing PAAR and rearrangement <u>h</u>ot<u>s</u>pot (Rhs) repeat motif found among effectors of the Rhs protein family [39], which is identical to Tre1 in its N-terminal region (amino acids 1–240). Because the protein codes a C-terminal Nicotine adenine dinucleotide (NAD) glycohydrolase domain (96.1% probability of similarity between its amino acids 261–402 and amino acids 35–207 of protein 4KT6_1 from *Streptococcus pyogenes*), we consider this protein an effector. A putative immunity protein encoding gene (GQQ23_RS09575) was located downstream. The protein is similar to the immunity protein Tsi6 (99.9% probability of similarity between amino acids 7– 100 and and the full length of Tsi6 of *Pseudomonas aeruginosa* (PDB: 4ZV0_B)). We hypothesize that these two proteins form a putative effector/immunity pair, because of the similarity that Tsi6 blocks NAD glycohydrolase activity of its effector [40].

The flexible effector/immunity island downstream of the *vgrG* island spans several genes whose functions we could not predict for all of them. Nevertheless, we located four effector/immunity pairs in this region of the T6SS cluster 1. The first pair is *rhsD/rhsI1*. The gene *rhsD* and its neighboring gene *rhsI1* were already discussed in *P. ananatis* as a putative effector/immunity gene pair [31]. In our set of strains, only strain A2 carries *rhsD* directly flanked by *rhsI1*, while *rhsI1* is absent strains R5, and T1 to T5 (Fig. 1B). The RhsD protein (a RhsA-family protein) carries a C-terminal Tne2-like effector domain having 97.3% probability of similarity between amino acids 1346–1460 and amino acids 15–131 of Tne2 (PDB: 6B12_A) according to HHpred. This Tne2 effector domain was identified in several RhsD homologs in different *Pantoea* species [41]. The immunity protein is similar to RhsI1 (PDB: 6XTD_B) from *Serratia marcescens* over its entire length (95.9% probability of similarity).

In the strains that lack *rhsD* (R1, B1, B3 and T6; Fig.1), we located a different putative RhsA-like protein (95% identity (81% coverage) to RhsD described above) together with a putative immunity protein encoding *rhsI* homolog. This RhsA/RhsI effector immunity gene pair is only present in strain R1. Strain B1 and B3 lack the *rhsI* gene, but harbor a copy of *rhsA*, while the strains T3, T4, T5 and T6 carry a pseudogene copy of *rhsA* that leaves the related *rhsI* as an orphan gene. The RhsI proteins of the two effector/immunity pairs, RhsD/RhsI1 and RhsA/RhsI, share only about 30% in sequence identity. Whether RhsD/RhsI1 and RhsA/RhsI are specific effector/immunity pairs, or whether they are redundant in function (which might explain why *rhsI* in strains T2, T3, T4, and T5 is not required), remains an open question.

The last two effector/immunity pairs that we identified in the 5’ region of the T6SS cluster 1 are the Pse2/Psi2 effector/immunity pair [29] in eight of the eleven strains and a putative Tre1/Tri1 effector/immunity pair present in only strain T6. The Tre1/Tri1 pair follows the canonical architecture observed in other T6SS systems with an upstream located cognate *vgrG* homolog (T6SS spike protein) followed by a specific Eag adaptor protein encoding gene [42,43].

Altogether, our analysis indicates the T6SS is functional in the examined *P. agglomerans* strains, as evidenced by the conservation of the core T6SS genes (i.e., *tssJ*– *tssC, tagH*–*tagE*; Fig. 1B). The heterogeneous content of effector and immunity pairs among the strains indicates that almost any pair of isolates is predicted to engage in some level of collateral killing.

### *Pantoea agglomerans* isolates harbor an active T6SS and show diverse levels of collateral inhibition and killing

For the experimental validation of T6SS function and the quantification of T6SS-mediated killing, we utilized *P. agglomerans* strain R1 [27] as the focal strain. The activity of the T6SS was validated by testing the ability of strain R1 to kill *Escherichia coli* MG1655 in direct competition. To enable direct selection of both competitors, strain R1 was equipped with a kanamycin-resistance encoding reporter plasmid (pTW1-mCherry; Supplementary Table S2), whereas *E. coli* MG1655 was equipped with a mini-Tn*7*::*dhfr*II insertion, conferring resistance against trimethoprim (Tm^R^). Competitions assays were performed by mixing both strains in equal frequencies followed by spotting the cell suspension on non-selective LB media plates. After four hours of incubation, the growth spots were harvested, serially diluted and plated on selective plates. When *E. coli* MG1655 was incubated with strain R1, we observed a severe reduction (10.000-fold) in the viable cell titer compared to controls without strain R1 (Fig. 2A). To validate that the growth inhibition of *E. coli* was caused by T6SS-mediated killing of strain R1, we generated a knock-out mutant of the T6SS Cluster 1 in strain R1 (hereafter R1 ΔT6SS, see Material and Methods). Performing head-to-head competitions between *E. coli* and R1 ΔT6SS showed no significant reduction in *E. coli* growth compared to axenic growth without R1 ΔT6SS (Fig. 2A). Taken together, our result demonstrates that *P. agglomerans* R1 harbors an active T6SS system and that T6SS-mediated killing under *in vitro* conditions depends on genes encoded in cluster 1.

**Figure 2.**
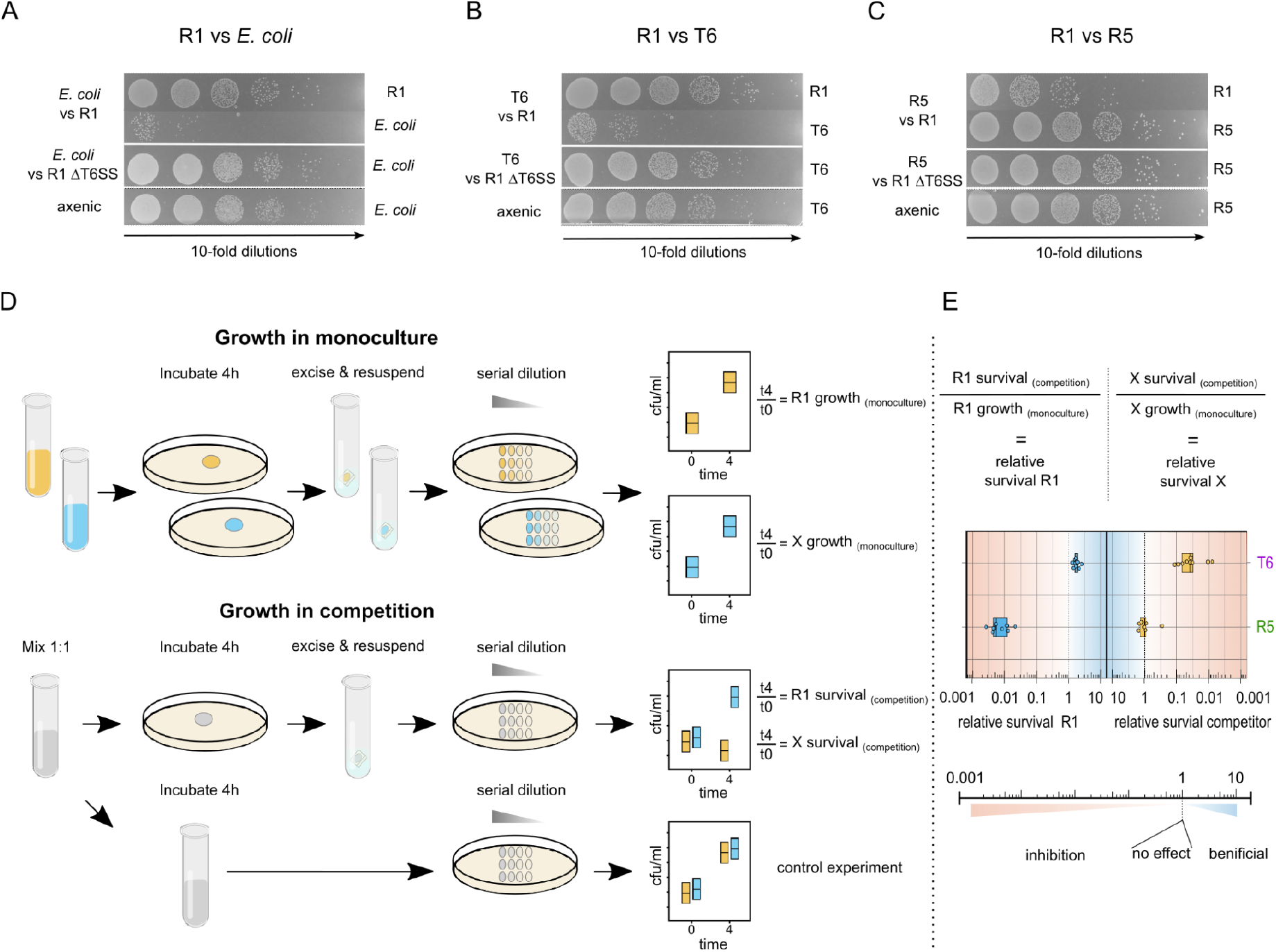
Validation of T6SS-mediated killing and competition pipeline. (**A**) Spotting assays of pairwise competitions between *P. agglomerans* R1 and *E. coli* MG1655, (**B**) *P. agglomerans* strain T6 and (**C**) strain R5. (**D**) Workflow for pairwise competition assays of strain R1 against the other ten strains in our strain set. The strains were cultured either mixed (1:1) or in monoculture on solid or in liquid medium. Population sizes (CFUs) were determined by plating at the onset of the experiment and after four hours of incubation. See Material and Methods for more detailed information. (**E**) Calculation of relative survival with example data from the competition of R1 against T6 and R5.

To compare the outcome of T6SS-mediated competition between distantly and closely related strains, we performed competition experiments between *P. agglomerans* R1 against the distantly related strain T6 and the closely related strain R5. To enable direct selection of the competitors after mixed incubation, both R5 and T6 were equipped with the reporter plasmid pRL153-gfp-cat encoding the green fluorescence protein (GFP) and the chloramphenicol resistance marker gene cat. Growth of T6 was ca. 1000-fold reduced in the competition with strain R1, but not in competition with the ΔT6SS mutant of R1 (Fig. 2B). Hence, the competition results are well explained by T6SS-mediated killing of T6 by R1. In contrast, strain R5 did not show growth deficiency in competition with strain R1, rather strain R1 displayed a slight (100-fold) reduction of growth when co-incubated with strain R5 (Fig. 2C). Altogether, our results show that T6SS-mediated killing is a common trait of *P. agglomerans* strains that can provide a competitive advantage against other bacteria under *in vitro* conditions.

### Quantification of T6SS-mediated competition

To assess the strength of T6SS-mediating killing among *P. agglomerans* strains we established an experimental pipeline to quantify the reciprocal inhibition potential of competing *Pantoea* strains (Fig. 2D). T6SS-mediated killing is known to be impaired during incubation in liquid media with shaking [44]. Therefore, all competition experiments were performed, in parallel, on solid and in liquid medium to ensure conditions for T6SS-mediated interactions/T6SS-permissive conditions compared to conditions that do not allow for T6SS-mediated interactions. Observing an inhibitory effect in competitions performed on solid medium, while no effect is seen in liquid medium, serves as confirmation of the effect of contact dependent killing. Cultures of the competing strains were grown either axenically or 1:1 in mixed culture (Fig. 2D). The relative survival of both strains post competition was calculated by dividing the ratio of the population sizes observed at t=4 hours and at the start of the experiment (t=0 hours) when grown in competition by the the ratio of the population sizes observed at the end (t=4 hours) and at the start of the experiment (t=0 hours) when grown in monoculture (Fig. 2E).

To test the approach performance and validate contact-dependent killing activity in *P. agglomerans* strains, we performed competition experiments of all eleven strains against *E. coli* MG1655. All *P. agglomerans* strains were equally effective against *E. coli* on solid medium, while no inhibitory effect on *E. coli* was observed after incubation in liquid medium (see Supplementary Fig. S3). The pairwise competition between strains R1 and T6 on solid media showed that relative survival of R1 was 1.74 ± 0.36 (median 1.66, n=9) and relative survival of T6 was 0.049 ± 0.034 (median 0.037, n=9) (Fig. 2E). In the competition between strains R1 and R5, relative survival of R1 was 0.0084 ± 0.0056 (median 0.054, n=9) and the relative survival of R5 was 1.05 ± 0.37 (median 1.01, n=9) (Fig. 2E). Thus, growth of R1 is strongly inhibited by R5 while the growth of R5 is not affected by the presence of R1. Notably, strain T6 and R5 showed both no difference in growth when grown with strain R1 in liquid medium as compared to growth in monoculture. Hence, our approach enables us to quantify the strength of reciprocal inhibition via a contact-dependent killing mechanism for both competing strains (Fig. 2E).

### Variation in T6SS-mediated killing corresponds to different effector/immunity gene repertoire

To explore the strength of competition under T6SS-permissive conditions, the pipeline described above (Fig. 2B) was used to compete our ten *P. agglomerans* strains against our focal strain R1. Head-to-head competition between strain R1 and the other strains resulted in either unidirectional or reciprocal killing. Notably, none of the strains exhibited mutual co-existence with R1 (Fig. 3B). The range of killing observed ranged from almost complete elimination of the competing strain, as observed in the competition between R1 and R5, to moderate levels of killing, as seen in the competitions between R1 and B1 or B3.

**Figure 3.**
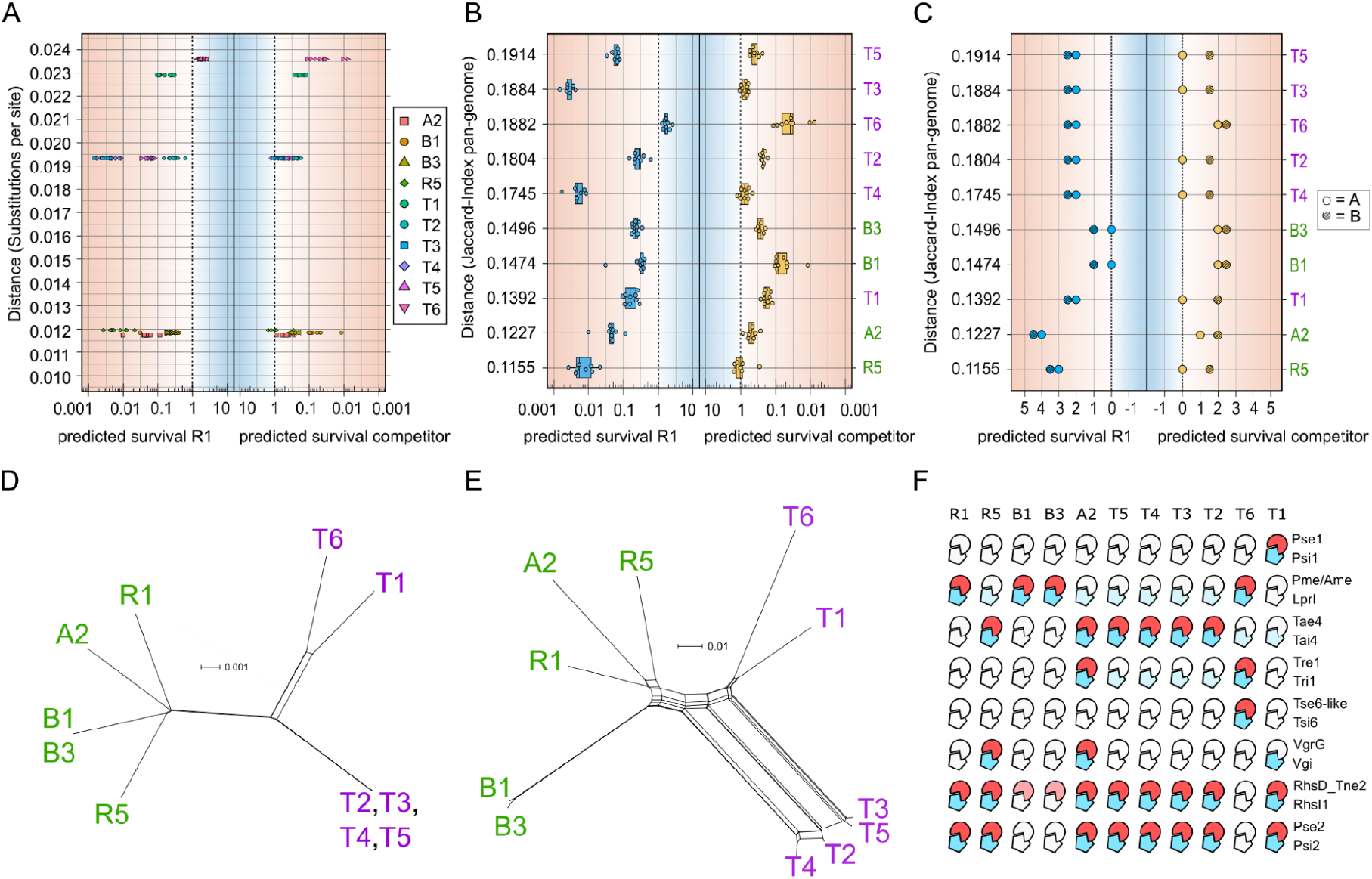
Relative survival in T6SS-mediated competition between related strains having different immune/effector genes. (**A**) Relative survival of strains in competition with strain R1 in relation to phylogenetic distance in substitutions per site (**B**) Survival of strains in competition in relation to phylogenetic distance based on pan-genome Jaccard distance (**C**) A prediction of the strength of collateral inhibition based on either differential effector and immunity genes (A) or both common and differential effector and immunity gene pairs (B). (**D**) Phylogenetic networks for the concatenated alignment of 3,157 complete single copy genes. (**E**) presence/absence pattern of 6,059 protein families across all the 11 strains. (**F**) Presence/absence panel of nine confirmed and predicted effector/immunity pairs in the 11 strains. Effectors are colored in red (light red for orphan effectors) and immunity proteins are shown in blue (light blue for orphan types). Absent effector/immunity factors are empty. Orphan effector proteins having distinctly different C-terminal domain compositions compared to the effectors whose cognate immunity protein was predicted, were excluded from the panel (i.e., VgrG in strains T5, T4, T3, T2).

To test if the observed range of relative survival was correlated with evolutionary distance, we first examined whether an association exists between the rate of survival and the number of substitutions per site between each strain pair. We found no significant correlation of relative survival and the number of substitutions per site for R1 (Spearman correlation p-value = 0.0785), as well as for the competitors (Spearman correlation p-value = 0.178; Fig. 3A). Complete single copy genes are characterized by a high level of conservation. This may hinder the detection of existing diversity across strains. In addition to single-nucleotide variants (SNVs), strain genomes vary in their gene content and differential gene gain and loss can result in different tree topologies than the ones inferred from SNVs. Indeed, we observe that relatively extant strains according to SNVs phylogenies are more closely related at the level of gene content (e.g., strains R1 and T1; Fig. 3D, E). To further investigate a possible association between T6SS-related killing effects and strain relatedness, we compared the relative survival to the Jaccard distance, which is based on the presence/absence of genes from the pangenome across strains. We found no significant correlation between the Jaccard distance and the strength of competition for R1 (Spearman correlation p-value = 0.0785), as well for the competitors (Spearman correlation p-value = 0.669). Altogether these results indicate that T6SS-mediated competition between strains is not associated with the phylogenetic relatedness neither at the level of SNVs nor at the level of gene content.

Previous studies suggest the presence/absence of effector/immunity pairs is the primary determinant for T6SS-mediated antagonistic interaction between closely related strains [19,45, and reviewed in reference 16]. Therefore, we next asked whether the level of competition depends on the repertoire of effector/immunity pairs of the strains. To examine whether the resulting competitions could have been predicted by the strains’ repertoire of effector/immunity pairs alone, we devised a scoring scheme based on common and differential effector/immunity pairs. In the simplest scoring scheme, the opponent inhibition score for a competing strain is calculated as the number of effector/immunity pairs not shared with the opponent. Nonetheless, the results of the pairwise competitions suggest that strains having the same effector/immunity pairs were capable of collateral inhibition (Fig. 3E). Therefore, we devised a second scoring scheme where the opponent inhibition score is increased by 0.5 for each pair of effector/immunity genes common for both competitors.

Comparing the prediction of the scoring schemes with the competitions, we found that both schemes correctly predict the dominant strain for each competition pair in almost all cases. The only exception being the competition between R1 and T6. Here, strain R1 clearly dominated T6 in competition, although their prediction did not overlap with the observed ranges of relative survival. The second scheme that accounts for identical effector/immunity pairs showed overall a slightly higher accuracy for the prediction relative survival (Fig. 3C, E). By comparing the repertoire of identified effector/immunity pairs of the eleven strains, we found that the dominant strain in competitions could be predicted solely by the presence/absence of Tae4/Tai4 and Pse2/Psi2. The results suggest a strong association between the inhibition of R1 and the presence of the Tae4 effector (A2, R5 and T2 to T5). In contrast, all strains dominated by R1 do not encode the Tae4 effector and do not possess Pse2/Psi2 (Fig. 3C, F). However, the calculated relative survival values from the competitions showed large deviations from the predicted outcome of both schemes. For example, strains T2 to T5 encode identical effector/immunity pairs, hence they were predicted to induce equally strong reduction of relative survival in R1 and receive a mild reduction in relative fitness by R1. In contrast to that strains T3 and T4 were among the strongest killers of R1, while T2 and T5 inflicted comparingly low levels of inhibition (Fig. 3C). Another example is strain A2, which inflicts less killing against R1 than strain R5, although it carries an additional effector/immunity pair, Tre1/Tri1, not present in R1 or R5 (Fig. 3C, F). Hence, the repertoire of effector/immunity pairs is a driving element in T6SS-mediated intraspecies competition, and our results indicate that additional factors influence the outcome of T6SS-mediated intraspecies competitions.

### Effector/immunity pairs are highly diverse across *Pantoea agglomerans* isolated from plants

To further study the evolution of T6SS components in naturally occurring strains, we compared the T6SS clusters among 113 publicly available *P. agglomerans* genomes mostly isolated from plants. To identify T6SS components across all studied genomes we searched for the presence of the corresponding homologs within protein families. Note that these families comprise protein sequences which are at least 30% identical over their global amino acids sequence. The core components of TSS6 cluster 1 were highly conserved across all studied isolates while the auxiliary components of the cluster show a variable presence and absence across the isolates (Fig. 4A). While, for example, the core components contractile sheath small subunit *tssB* and tube protein *hcp* genes have been found in all the studied isolates while, the gene for the *pse1* effector is found in almost half of the strains, the gene for the immunity gene *psi1* is found in only a few strains (i.e., *pse1* is found in 53 strains and *psi1* is found in 8 strains. Fig. 4B). The variable region includes the effector/immunity pairs we identified here (Fig. 1B; Fig. 3F), and additional hypothetical proteins that may correspond to yet unidentified effector/immunity genes. The presence and absence pattern of protein families in the variable region may stem from differential gene gain and loss. It has been previously suggested that genetic recombination is frequent in the evolution of T6SS (e.g., in *Vibrio cholerae* [17,46]). The mosaicity observed further suggests an accelerated evolution of the genomic loci that harbor the effector-immunity pairs.

**Figure 4.**
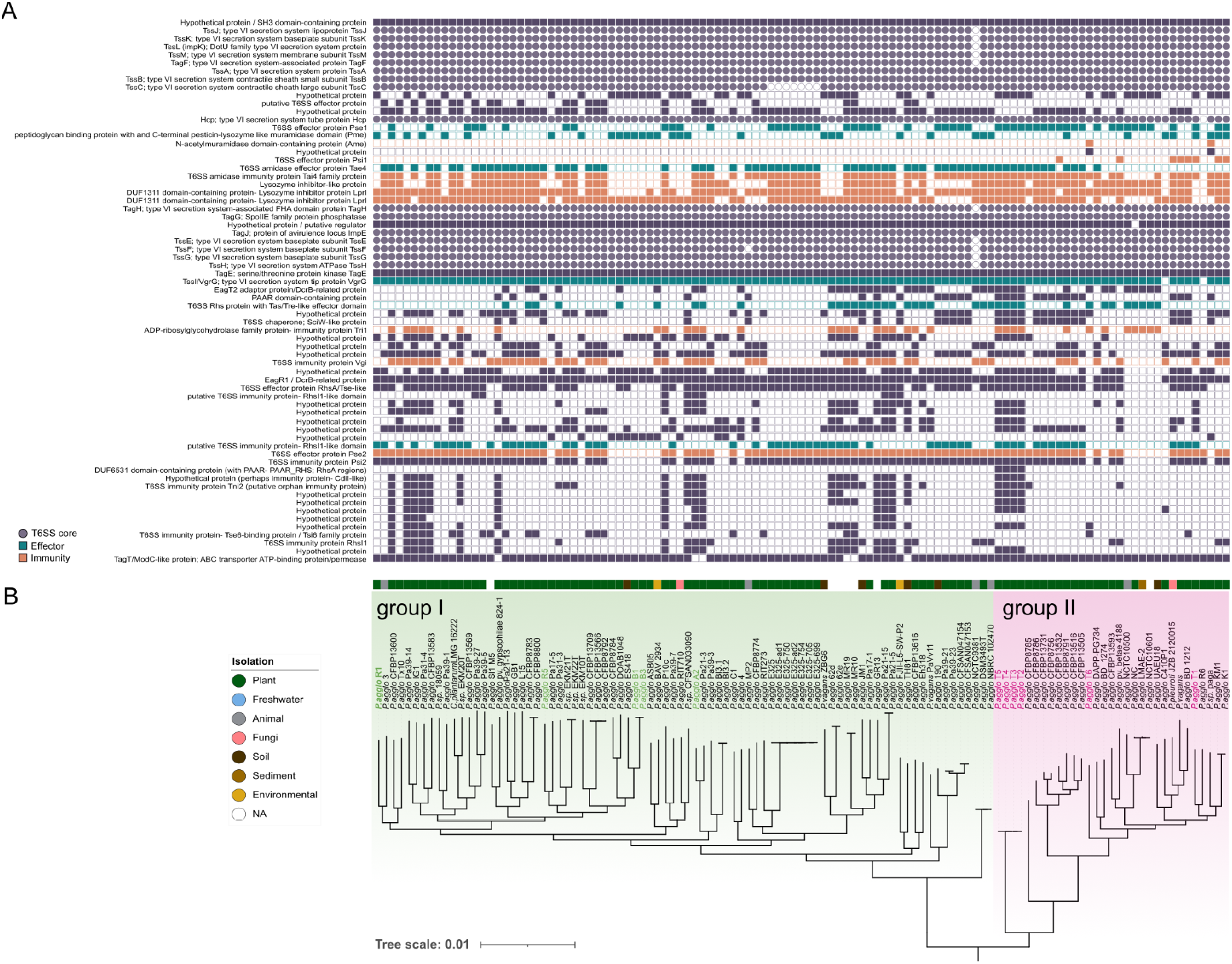
The repertoire of T6SS genes and phylogeny of *P. agglomerans* isolates. (**A**) Gene presence/absence pattern of 69 protein families in the T6SS cluster 1 in *P. agglomerans* across 113 genomes. Core components of the T6SS are represented as circles while immunity and effector proteins are highlighted in orange and blue, respectively. (**B**) A phylogenetic tree of the isolates was inferred from 880 single-copy universal genes. The branch length shows the number of substitutions per site. The root position inferred with MAD [47]. Isolates from *T. aestivum* cultivars Akteur, Benchmark and Runal are shown in green. Isolates from landrace *T. aestivum* are shown in pink. Group I and group II are enclosed in green and pink boxes, respectively.

To study the evolution of cluster-1 genes, we first inferred the phylogenetic species tree of the aforementioned strains. The rooted phylogenetic tree reveals a clear split between *P. agglomerans* isolated from seeds of wheat cultivars (Runal, Akteur & Benchmark) and those isolated from seeds of wheat landrace (Fig. 4B) [27]. In the following, we will refer to the groups defined by the root of the species tree as group I and group II strains (Fig. 4B).

The absence of a clear association between phylogenetic relatedness and the strength of competitive inhibition may be explained by horizontal acquisition of effector-immunity proteins (via genetic recombination or horizontal gene transfer). Indeed, the pairwise proportion of common effector-immunity genes and the total shared gene content between strain pairs are only weakly correlated (r_s_=0.11, p-value < 2.2×10-16, using Spearman, both measures calculated with Jaccard index). Nevertheless, the phylogenetic distance between *P. agglomerans* strains, expressed as the number of substitutions per site, is significantly correlated with the difference in gene content (r_s_=0.45, p-value < 2.2×10-16, using Spearman). This result suggests that effector/immunity pairs have a different mode of evolution compared to the rest of the *P. agglomerans* pangenome. To test for differences in evolutionary rate between the core T6SS genes and effector/immunity pairs, we compared core and effector-immunity gene trees. The results show that gene trees of effector/immunity pairs have a significantly larger median branch length (Wilcoxon rank sum test p-value = 0.023. Supplementary Fig. S2).

Altogether, these results suggest that effector/immunity pairs evolve at a higher pace in comparison to core T6SS in cluster 1, which may contribute to the diversity of gene presence/absence in the variable region. Previous studies suggested that secreted proteins undergo an accelerated rate of evolution and may experience intra-chromosomal gene conversion; these evolutionary scenarios may explain why secreted proteins often form multigene families (i.e., where organisms harbor several gene duplicates in the genome [48]. Here, we find that six out of eighteen effector-immunity gene families and 9 out of eighteen core T6SS component gene families are multigenic. In our dataset, families of several effector and immunity proteins are multigenic, including *vgrG, rhsA/rhsD, pse1, tri1, tre1* and *pme*, indicating a role of gene duplication or gene acquisition in their evolutionary history.

Previous studies suggested that the evolution of type III secretion system effectors in pathogenic bacteria is accompanied by intra-chromosomal rearrangements leading to ‘terminal reassortment’, where gene fission and fusion give rise to the origin of new effectors [49]. To study if intra-chromosomal rearrangements may also be involved in the evolution of the T6SS cluster 1 in *P. agglomerans*, we searched for intra-chromosomal sequence similarity within the T6SS cluster 1 in the 11 studied isolates. Indeed, duplication events are rampant in that locus, with different outcomes for the effector-immunity gene content in the different strains (Fig. 5A). For example, in strain A2, three regions within cluster 1 have been duplicated. The first duplicated region involves the *vgrG* and *vgrG* effector genes. A similar duplication is observed in several strains (R5, A2, T5 and T6), where it indicates that the *vgrG* effector and core *vgrG* component have a common origin by duplication, which likely occurred in the ancestor of the strains. In T6, this duplication spans additionally two other genes, one of which is *tre1* that is found duplicated in a region giving rise to *tri1 A* and *tri1 B* copies. The second duplicated region in A2, involves an *rhsD* effector gene and a fragment in a non-coding region downstream of this gene. This region appears duplicated twice in T5, as part of a pseudogene and an annotated functional gene, both characterized by the presence of Rhs domains. Notably, *rhs* domains are known for their implication in genome rearrangements and domain reshuffling in *Escherichia coli* [50]. We hypothesize that the original duplication was already present in the ancestor and underwent different degrees or erosion in the lineages leading to the two strains. A third region is duplicated twice and involves three *tri* immunity genes. Two of them are identified as paired (*tri1* A and *tri1* B) and a third one (*tri1*) is identified as an orphan immunity gene. In T5, the *tri1* A copy allocated within the *vgrG* island is absent. The observed pattern of sequence similarity indicates the occurrence of intra-chromosomal recombination in the variable region of cluster 1.

**Figure 5.**
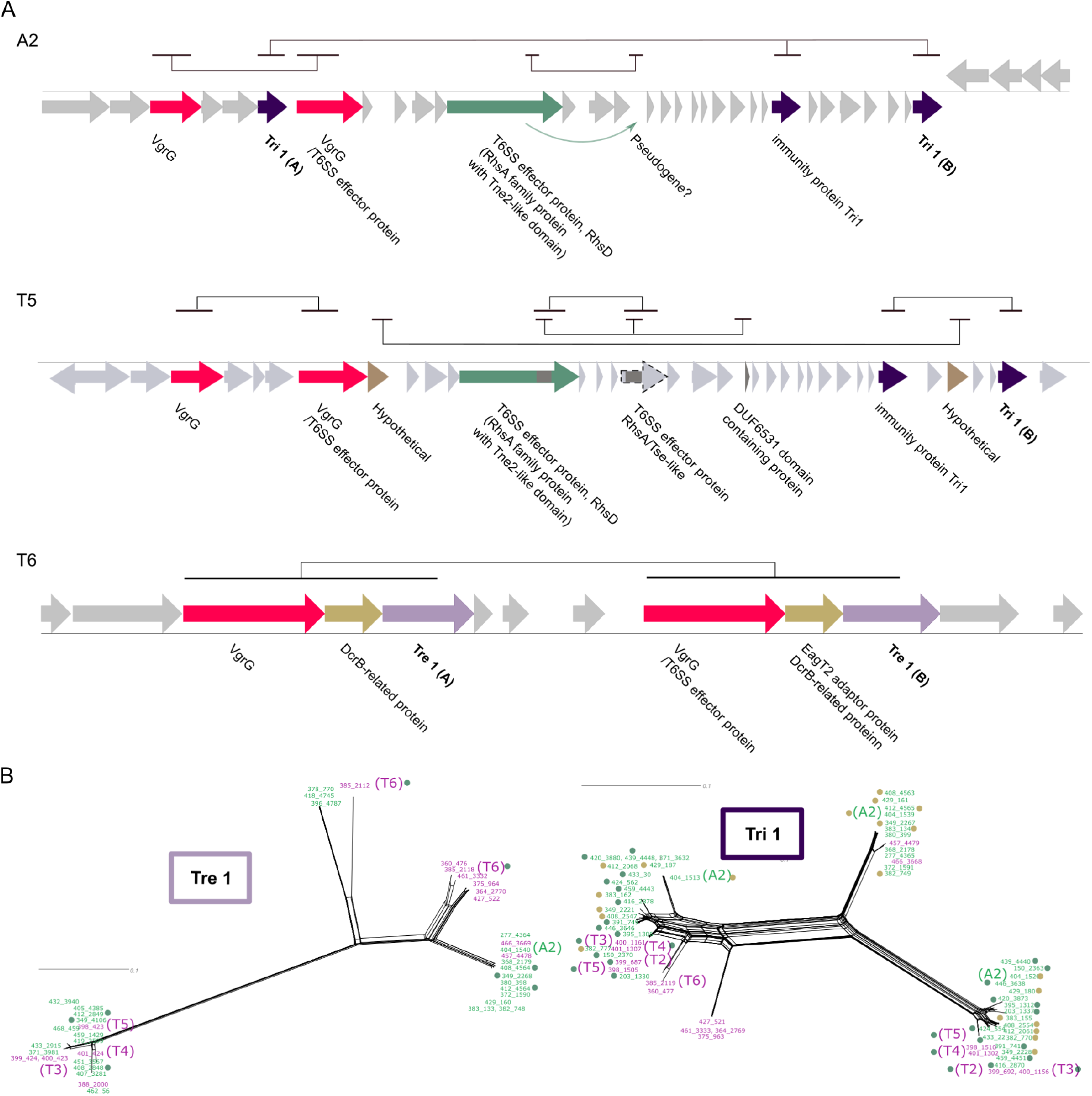
Intra-chromosomal T6SS homologous regions and effector/immunity protein phylogeny. (**A**) Intra-chromosomal sequence similarity detected with gepard [51] for three selected strains. Open reading frames (ORFs) in these regions are depicted by arrows. Pseudogenes are indicated with dashed outer lines. Lines above the ORFs diagram indicate the detected duplications. Genes belonging to the same gene family are shown in the same color. ORFs outside duplicated regions are shown in gray. Bold names correspond to predicted effector/immunity pairs. (**B**) Phylogenetic network reconstructed for the *tri1-tre1* pair. Sequences from isolate groups I and II are highlighted in green and pink, respectively. Yellow and green circles next to the OTUs indicate that the corresponding strain harbors three and two copies of the gene, respectively. Strains with no circle harbor a single gene copy.

To further explore evidence for gene duplication, we reconstructed the phylogeny of effector and immunity gene protein families. The phylogenetic network reconstruction reveals a clear divergence of multiple gene copies in the same genome, thus the duplication events are unlikely to be recent. For example, *tre1*, which is present in two copies in strain T6, is found in two different splits of the phylogeny (Fig. 5B). Although no other strain from the initial set of eleven isolates harbors a copy of this gene, we observe that the presence of multiple *tre1* copies is not specific to T6, and can be observed in three additional strains (Fig. 5B). The *tre1* phylogeny further reveals one variant specific to group II isolates. Such a pattern is more likely to emerge following gene duplication rather than independent gene transfers. The *tre1* phylogenetic network suggests that the ancestor of the entire species harbored three *tre1* paralogs, where differential gene loss, likely by intra-genomic recombination in Rhs regions, played a role in the evolution of that effector protein family. The phylogeny of the immunity protein paired with *tre1* - *tri1* - reveals a larger number of strains carrying multiple copies of the immunity gene; indeed, 20% of the strains harbor three *tri1* copies, most strains harbor two copies (43%) and the remaining 37% strains encode a single copy. The *tri1* copy number does not seem to correspond to the species phylogeny, indicating multiple gene duplications (and loss). Taken together, the phylogenetic reconstruction supports the inference of gene duplication events that can be traced to both old and recent intra-chromosomal recombination events in the *P. agglomerans* isolates.

## Discussion

A growing number of reports document how the T6SS modulates and shapes the assembly of host-associated microbiota in various host systems (reviewed in [52] and [53]). For any bacterial cell, the benefit of a functional T6SS system manifests in a potent mechanism for competitive exclusion of other bacterial cells being vulnerable to translocated effectors.

In this study, we investigated the outcome of intraspecific competition in a set of eleven closely related environmental isolates of *P. agglomerans* characterized by a limited but increasing phylogenetic distance to each other. We find that all eleven strains harbor up to three distinct T6SS-related gene clusters in their chromosomes. In contrast to other plant-associated enterobacterial species [54], no T6SS-related gene clusters were detected on the genus-specific plasmids, neither on the genus-specific large *Pantoea* plasmid (LPP) nor the second large plasmid that is characteristic for *P. agglomerans*. Indeed, all eleven strains effectively kill vulnerable *E. coli* cells lacking the immunity profile to any of the attacking strains. Killing is abolished when the genes encoding the main structural components are deleted from Cluster 1, which proves that T6SS-mediated killing does not require components encoded in Cluster 2 and 3. Assessing the architecture of the T6SS Cluster 1 in the eleven isolates revealed distinctive genetic differences in the individual effector/immunity gene repertoires. Our analysis shows that despite a heterogeneous distribution of known and putative effector/immunity gene pairs across the inspected strains, every strain harbors a functional T6SS that is predicted to engage in some level of collateral killing.

To assess the effects of T6SS-mediated competition on the intraspecific level, we developed an *in vitro* experimental pipeline for the quantification of T6SS-mediated killing. Using this approach we show that interbacterial killing occurs in varying degrees between these phylogenetically closely related strains. Using a star-shaped investigation scheme, we quantified bidirectional T6SS-mediated killing interactions relative to our focal strain R1, instead of competing all strains in a pair-wise manner against each other. This approach together with the analysis of the effector/immunity repertoire of each strain, however, is sufficient to show that the general outcome and effectiveness of T6SS-mediated collateral killing between closely related strains is not associated with phylogenetic relatedness/distance. In other words, mutually compatible strains are required to have a close-to-identical degree of relatedness within the cluster in order to coexist, albeit still in competition for common resources.

In line with this view and despite limited success in assigning putative functions to genes encoding the numerous hypothetical proteins found in strain A2 and the strains T2 to T5, the observed patterns of collateral killing are qualitatively correct reflected in their overall direction when using a simple scoring scheme based on a presence/absence matrix of the effector/immunity pairs that we identified.

Although our experimental scheme was not intended to identify mutually compatible strains that would be immune to each other’s attack, an analysis of the effector/immunity gene pairs in the strains suggests that the interaction between strain B1 and B3, as well as between the strains T2 to T5, is likely to be neutral. These strains have an identical repertoire of effector/immunity gene pairs, which means that their T6SS profiles would be compatible and recognize each other as self. However, recent studies have described kin discrimination systems that rely on subtle differences within conserved effector/immunity pairs in different bacterial species, including *Proteus mirabilis* [55,56] and *Vibrio cholerae* [57]. For example, even slight differences in one immunity protein can cause related strains carrying an otherwise similar effector/immunity pair repertoire to lose immunity against each other’s effector.

To test for such subtle differences, we looked at the sequence similarity level of relevant effector or immunity proteins that may have played a role in some of the interactions we observed. While the closely related strains T2, T3, T4, and T5 all displayed relatively high survival rates when in competition with strain R1, their impact on strain R1 did not match our model’s prediction. Because cluster 3 is absent in these four strains, differences in their effector/immunity pairs located within cluster 1 would provide an explanation for the stark differences in the relative survival of strain R1. Although the four strains are fully syntenic across the entire cluster 1 and encode identical effector/immunity pairs, differences in their effector/immunity pairs located within cluster 1 could explain the stark differences in the relative survival of strain R1. A notable difference is the presence of a *vgrG* homolog of the specialized VgrG-effector present in strain A2. This VgrG protein in the four strains lacks the effector domain and hence, a toxic effect on R1 is not plausible. However, we found no differences in the putative RhsD effector proteins or the Tae4 effector protein, which is identical on the protein level in the strains T2 to T5. We also found that the putative RhsI1 immunity protein in strain A2 shares 75% identity with the RhsI protein present in the other strains (i.e., R1, R5, and T2 to T5), where it is fully conserved across strains. This difference, however, does not explain the differences in the relative survival of strain R1, indicating that other factors are also affecting relative survival beyond the identified effector/immunity pairs.

The prevalence of T6SS clusters in plant-associated *P*.*agglomerans* strains shows that T6SS-mediated interactions is a hallmark of that species. The presence of homologous fragments within the T6SS cluster 1 region involves effector and immunity proteins as well as Rhs-domain containing proteins. This suggests that intra-chromosomal recombination likely plays a crucial role in the origin and evolution of effector/immunity pairs in *P. agglomerans*. Such mode of evolution has been previously described for the type IV secretion system in the pathogen *Bartonella*, where the effectors are shown to originate by duplication and diversification [58]. Here, we suggest that similar mechanisms are involved in the evolution of effector/immunity pairs in the type VI secretion systems as well. Frequent recombination and differential loss of effector/immunity genes serves as an explanation why the killing capabilities of the strains are not associated with phylogenetic relatedness.

On the level of intra-species interactions of host-associated bacteria, competition during colonization of an emerging host-habitat like a plant seedling, phylogenetically closely-related strains with similar metabolic capabilities are expected to compete fiercely when attempting to settle in the same niche. Thus, T6SS-mediated competitive exclusion may have a stark effect on the assembly and diversity of the host microbiota. Most T6SS related studies in the field of plant associated bacteria focus on inter-species competition, thus the role of T6SS-mediated intra-species competition remains understudied. In this study we showed that closely related *P. agglomerans* strains indeed compete fiercely in a T6SS dependent manner. T6SS-mediated interactions may lead to competitive exclusion resulting in pronounced monopolization effects in the plant host habitat. Monopolization effects are likely to be more pronounced during early stages of host development, while the carrying capacity and size of the host habitat are limited as recently shown in the Squid-*Vibrio* model [59]. Outcomes of such intra-species competitions likely depend on the diversity of the bacterial founder population which is associated with the mode of transmission of the bacterial strain. Low diversity in the founder population, associated with vertical transmission, is seemingly more likely to result in a monopoly for a single strain in a given niche. Horizontal transmission, accompanied with a higher diversity of closely related strains, hence it is more likely to result in a mixed assembly of strains. During later stages of host development T6SS-mediated intra-species competition may lead either to the replacement of a closely related strain or to the generation of sectors of functionally redundant, but genetically diverged strains. Future *in planta* experiments will help to reveal how T6SS-mediated competitions between closely related strains affect the microbiome assembly and diversity within the plant habitat.

## Methods

### Strains and cultivation

*P. agglomerans* and *E. coli* were grown in lysogeny broth (LB) supplemented with either kanamycin, 10 µg/ml; chloramphenicol 10 µg/ml, trimethoprim 50 µg/ml, ampicillin 100 µg/ml or 25 µg/ml spectinomycin as appropriate. All cultures were grown at 30°C if not indicated otherwise. The RSF1010-derived reporter plasmids encoding either GFP or mCherry were introduced into *P. agglomerans* by electroporation. Bacterial strains and strain/reporter plasmid combinations used in this study are provided in Supplementary Table S2.

### Recombinant DNA techniques

Molecular cloning techniques used in the construction of plasmid were done by using standard procedures. The plasmids for this work were constructed using the Gibson-Assembly technique (NEBuilder protocol, New England Biolabs, Germany). Plasmid constructions were first established in *E. coli* pir1 (Thermo Scientific, Germany). Plasmid-DNA isolations were performed by alkaline lysis. Genomic DNA was isolated from *P. agglomerans* R1 using the Wizard Genomic DNA Purification Kit (Promega, Germany). All restriction enzymes were obtained from New England Biolabs GmBH and used according to manufacturer’s protocols. Phusion^™^ High-Fidelity DNA-Polymerase (Thermo Scientific, Germany) was used in all PCR-based cloning steps. PCR amplicons and restriction digestions were purified using the GeneJET Gel Extraction and DNA Cleanup Micro Kit (Thermo Scientific, Germany). All plasmids constructed in this work were analyzed by restriction endonuclease digestion and Sanger DNA sequencing.

### Generation of *P. agglomerans* R1 mutant strains

The deletion of the T6SS Cluster 1 was performed using a λ-Red-mediated allele replacement strategy as described by [60]. The cluster was replaced with a *sacB*-*nptII* cassette. Primer used for the amplification of the chromosomal regions flanking the cluster 1 are given in Supplementary Table S3. The two PCR fragments were cloned into the gene targeting vector pGT42 [61]. The resulting plasmid served as template for the PCR amplification of the (*sacB*-*nptII*)-cassette embedded into the natural flanking regions of the T6SS cluster 1. To allow for λ-Red-mediated integration, the (*sacB-nptII*)-cassette was transformed into cells of strain R1 carrying pTKRED. Electrocompetent cells of *P. agglomerans* R1 carrying pTKRED were prepared by diluting cells of an overnight culture 100-fold in fresh LB supplemented with 25 µg/ml spectinomycin. After one hour of growth at 30°C with shaking, the culture was amended with 6 mM isopropyl-ß-D-galactopyranoside (IPTG). Once the culture reached an OD_600_ of 0.5, cells were harvested by centrifugation and washed twice in 10% glycerol and concentrated as described [61]. Electrocompetent cells were transformed with 1 µg of the purified PCR fragment (parameters: 2,5 kV, 25 µF, 200 ?, cuvette with 1 mm gap). Transformants were selected on LB plates supplemented with 10 µg/ml kanamycin. Selected clones were cured from pTKRED by streaking on non-selective LB agar and incubation at 37°C overnight. The loss of pTKRED was validated by streaking on spectinomycin-supplemented LB plates and by PCR. Allelic replacement of T6SS cluster 1 was confirmed by PCR and Sanger sequencing.

### Competition-Assays

Overnight cultures of competing strains were grown separately overnight in LB medium with the appropriate antibiotic at 30°C. For a list of strains see Supplementary Table S2. The cultures were normalized to an OD_600_ of 0.5. For each competition, the OD-adjusted cultures of the two competition partners were mixed in a 1:1 ratio and 20 µl of the mixture was spotted on non-selective LB-plates and incubated at 30°C. For quantification of growth of the individual strains in monoculture, 10 µl of OD-adjusted cultures were spotted on LB-plates and incubated as described above. Viable cell titers were quantified at the onset of the incubation (t=0 h) and after four hours (t=4 h). Cells were serially diluted and plated on non-selective LB plates and, in case when separate selection for the respective competitor genotypes was required, plated on LB plates supplemented with the appropriate antibiotic. After 4 hours of incubation, the cells grown in competition or monoculture were harvested from the plates and resuspended in one ml PBS prior to plating as described above. In parallel to the incubation on solid surface, 20 µl of the mixed competitor cultures as well as 10 µl the monocultures were used for inoculation of one ml LB medium and incubated with shaking at 30°C for four hours and processed as described above. Each experiment was repeated three times with three replicates each. The relative fitness of a strain was calculated by dividing the ratio of CFUs at t = 4 h and t = 0 h with the ratio of CFUs from the respective monoculture (t = 4 h and t = 0 h; see Fig. 2B).

### Bioinformatics

*Pantoea* genomes were retrieved from NCBI in March, 2021 and used for comparative analysis. Protein families were reconstructed by extracting reciprocal best blast hits among all the proteins with an e-value < 1×10^−6^ [63]. Global pairwise alignments were then calculated with parasail using Blosum62 as amino-acid substitution matrix [64] and only hits with a minimum sequence identity of 30% were retained. Protein were clustered into families using the MCL algorithm [65] with an inflation index of 2. For this study, we considered the genomes of 101 *P. agglomerans* and 12 additional isolates, which have been putatively missanotated as different *Pantoea* species based on the phylogenetic reconstruction (Fig 4B). Phylogenetic reconstruction of the species tree was performed using a concatenated alignment of 880 single-copy universal genes found across all isolates. Single gene and protein alignments were constructed with MAFFT using the default parameters [66]. The phylogeny for the full *P. agglomerans* dataset was inferred with IQTREE version 2.2 [67] using a general evolutionary model GTR+F, a partitioned analysis and 1,000 bootstrap replicates. Likewise, a phylogenetic tree was inferred for 11 selected isolates using 3,157 single-copy universal genes shared among the selected genomes. Pairwise phylogenetic distances were extracted from the tree using the R package ape [68]. Pairwise pangenome Jaccard distances were calculated using the presence of the 6,059 and 13,926 proteins that comprise the pangenome of the 11 and selected 113 total isolates. Similarly, the T6SS Jaccard distance was calculated based on the presence of proteins which were identified as T6SS-Components (see below). Jaccard distances were calculated with the R package vegan (https://CRAN.R-project.org/package=vegan). Phylogenetic networks were inferred with SplitsTree [69] using the concatenated alignment of single-copy universal genes and the concatenated binary code for the presence of genes in the pangenome and the T6SS, respectively. Additionally, single protein family phylogenetic networks were inferred using multiple sequence alignments of the protein sequences. Intra-chromosomal similarity was inferred from dotplots created with gepard [51].

### Re-annotation of T6SS-Components

The positions of the T6SS gene clusters in our *Pantoea* strains were first determined based on the available genome annotation information of the genomes already automatically annotated by the NCBI. Individual genes in the clusters were identified by homology searches using the protein sequences of T6SS-associated genes that were previously reported in other *Pantoea* strains. The analysis of hypothetical proteins and uncharacterized genes lacking an annotation was performed manually via homology searches using the data set of experimentally validated T6SS components available from the SecReT6 database [70] and remote protein homology detection using hidden Markov modeling (HHpred; [35]).

