## Supplementary Tables and Figures for "T6SS-mediated competitive exclusion among *Pantoea agglomerans* associated with plants"

#### Supplementary Tables S2 – S3

#### Supplementary Figures S1 – S4

### T6SS-mediated competitive exclusion among *Pantoea agglomerans* associated with plants

Kai Ripcke[1], Devani Romero Picazo[1], Shreya Vichare[1], Daniel Unterweger[2,3], Tal Dagan[1], Nils F. Hülter[1]

[1]Institute for General Microbiology, Kiel University, 24118, Kiel, Germany.

[2]Institute for Experimental Medicine, Kiel University, Michaelisstraße 5, 24105 Kiel, Germany.

[3]Max Planck Institute for Evolutionary Biology, August-Thienemann-Straße 2, 24306 Plön, Germany.

**Supplementary Table S2:** Bacterial strains and Plasmid used in this study.

| Strain or Plasmid | Characteristics[a] | Source |
|---|---|---|
| **Strains** | | |
| ***Escherichia coli*** | | |
| MG1655 | wild-type | Thermo Scientific |
| MG1655 pRL153-GFP | Derivative of *E. coli* MG1655, transformed with pRL153-GFP, $Km^R$ | Wein et al. 2018 |
| MG1655 mini-Tn7::*dhfrII* | mini-Tn7::*dhfrII,* $Tm^R$ | Wein et al. 2018 |
| pir1 | F-Δlac169 rpoS(Am) robA1 creC510 hsdR514 endA recA1 uidA(ΔMluI)::pir-116 | Thermo Scientific |
| ***P. agglomerans*** | | |
| R1 | Wild-type | Soluch et al. 2021 |
| R1 pTW1-mCherry | Derivative of *P. agglomerans* R1, transformed with pTW1-mCherry, $Km^R$ | Soluch et al. 2021 |
| R1 pTKRED | Derivative of *P. agglomerans* R1, transformed with pTKRED, $Sp^R$ | This study |
| R1 ΔT6SS-1 | Derivative of *P. agglomerans* R1 T6SS-1 cluster deletion, $Km^R$ | This study |
| R5 | wild-type | Soluch et al. 2021 |
| R5 pRL153-cat-GFP | Derivative of *P. agglomerans* R5 transformed with pRL153-cat-GFP, $Cm^R$ | This study |
| A2 | wild-type | Soluch et al. 2021 |
| B1 | wild-type | Soluch et al. 2021 |
| B2 | wild-type | Soluch et al. 2021 |
| T1 | wild-type | Soluch et al. 2021 |
| T2 | wild-type | Soluch et al. 2021 |
| T3 | wild-type | Soluch et al. 2021 |
| T4 | wild-type | Soluch et al. 2021 |
| T5 | wild-type | Soluch et al. 2021 |
| T6 | wild-type | Soluch et al. 2021 |
| T6 pRL153-cat-GFP | Derivative of *P. agglomerans* T6 transformed with pRL153-cat-GFP, $Cm^R$ | This study |
| **Plasmids** | | |
| pTW1-mCherry | RSF1010 replicon, fluorescence protein mCherry, $Km^R$ | Wein et al. 2018 |
| pRL153-GFP | RSF1010 replicon, fluorescence protein GFPmut3.1, $Km^R$ | Tolonen et al. 2006 |
| pRL153-cat-GFP | Derivative of pRL153-GFP, $Cm^R$ | This study |
| pTKRED | phage λ *gam, bet* and *exo* genes under control of $P_{lacUV5}$, | Kuhlman & Cox 2010 |
| pGT42 | CmR, *SacB* | Wein et al. 2018 |
| pGT42-T6SS-1-$_{sub}$ | CmR, $Km^R$, $Amp^R$ (nptII ($Km^R$)-*sacB* cassette surrounded by a 408 bp upstream and 450 bp downstream sequence homologous to *P.agglomerans* R1 chromosome up- and -downstream of T6SS-Cluster1 | This study |

[a] $Amp^R$, $Cm^R$, $Km^R$, $Sp^R$, and $Tm^R$ = resistance against chloramphenicol, kanamycin, spectinomycin, and trimethoprim, respectively.

**Supplementary Table S3**. List of primers used in this study. Presented are their sequence, Tm and usage used in this study.

| Primer name | Primer sequence 3´-5´ | Tm (°C) | Usage |
|---|---|---|---|
| T6SS-KO-up-fw | GCCTCACTGATTAAGCACTGGTAACTT CTGTAAACTCGCAGCTTATG | 85 | Amplification of a homologous Fragment upstream of T6SS-Cluster 1 in *P. agg* R1 |
| TSS6-KO-up-rv | CTCTTTGTCGTATCCACGTTCTAGAGTT AAAGTGCCATGGCGAGG | 83 | Amplification of a homologous Fragment upstream of T6SS-Cluster 1 in *P. agg* R1 |
| TSS6-KO-down-fw | ATCGCCTTCTTGACGAGTTCTTCTAGAGT TTCCGTAATGCTGGCAG | 83 | Amplification of a homologous Fragment downstream of T6SS-Cluster 1 in *P. agg* R1 |
| TSS6-KO-down-rv | CAAAAGCTGGAGATCAAGTTCGAGTAC AATCCATTGCTGC | 82 | Amplification of a homologous Fragment downstream of T6SS-Cluster 1 in *P. agg* R1 |
| T6SS-seq-up | GCCACGTATTACGGTCACC | 76 | Sequencing of T6SS-Cluster1 knock-out in *P.agg* R1 |
| T6SS-seq-down | CCGAACAAGATAACGCTGTGC | 79 | Sequencing of T6SS-Cluster1 knock-out in *P.agg* R1 |
| bla-3´-seq | GTCTCGCGGTATCATTGCAG | 58 | Sequencing of pGT42-T6SS-Cluster 1-KO |
| sacB-p-sq | CAGCAGTGCGGTAGTAAAGG | 59 | Sequencing of pGT42-T6SS-Cluster 1-KO |
| nptII-3´rev | GCTCAGAAGAACTCGTCAAGAAGG | 49 | Sequencing of pGT42-T6SS-Cluster 1-KO |
| NH160 | GCCAGGCGCGCcATTAACCCTCAC | 54 | Sequencing of pGT42-T6SS-Cluster 1-KO |

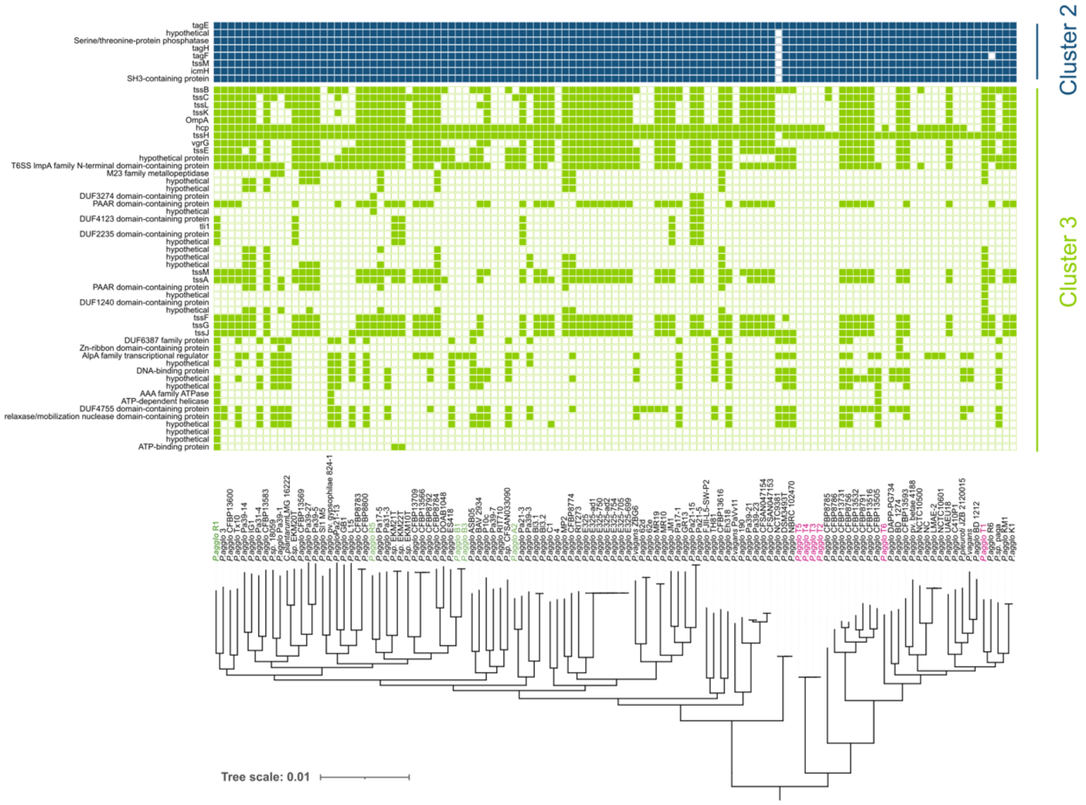


**Supplementary Figure S1.** The repertoire of cluster 2 and 3 T6SS genes and phylogeny of *P. agglomerans* isolates. (A) Gene presence/absence pattern of 8 and 48 protein families in the T6SS cluster 2 and 3, respectively in *P. agglomerans* across 113 genomes. The phylogenetic tree of the isolates was inferred from 880 single-copy universal genes. The branch length shows the number of substitutions per site. The root position inferred with MAD (Tria et al., 2019). Isolates from *T. aestivum* cultivars Aktuer, Benchmark and Runal are shown in green. Isolated from landrace *T. aestivum* are shown in pink. group I and group II are enclosed in green and pink boxes, respectively. T6SS cluster 2 shows an extreme level of conservation across the entire region while T6SS cluster 3 shows even higher mosaicity than cluster 1 with only two genes (hcp and tssH) present in most of the strains. The lack of conservation among its genes further supports the lack of functionality in T6SS cluster 3.

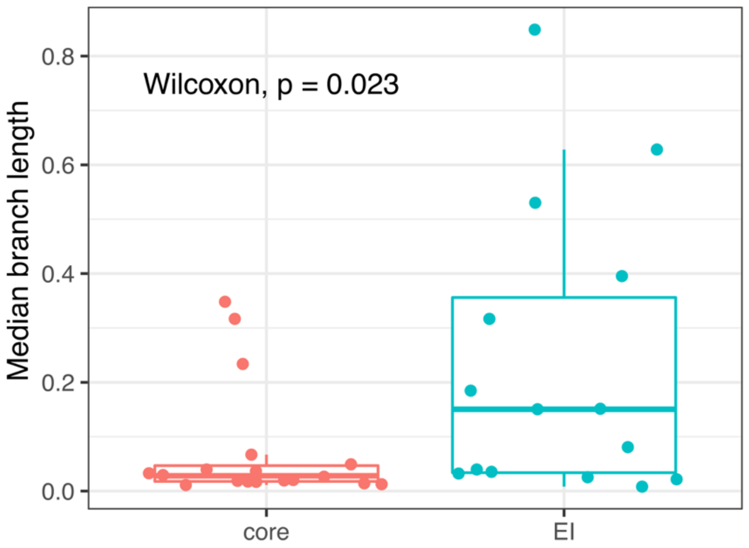


**Supplementary Figure S2**. Average branch length of T6SS gene families. Median branch length among pairs of strains estimated as the median substitution per site extracted from gene family phylogenies for core T6SS components and E-I pairs. Median branch length core = 0.028. IQR branch length core = 0.0291. Median pairwise branch length E-I pairs = 0.150. IQR branch length E-I pairs = 0.322.

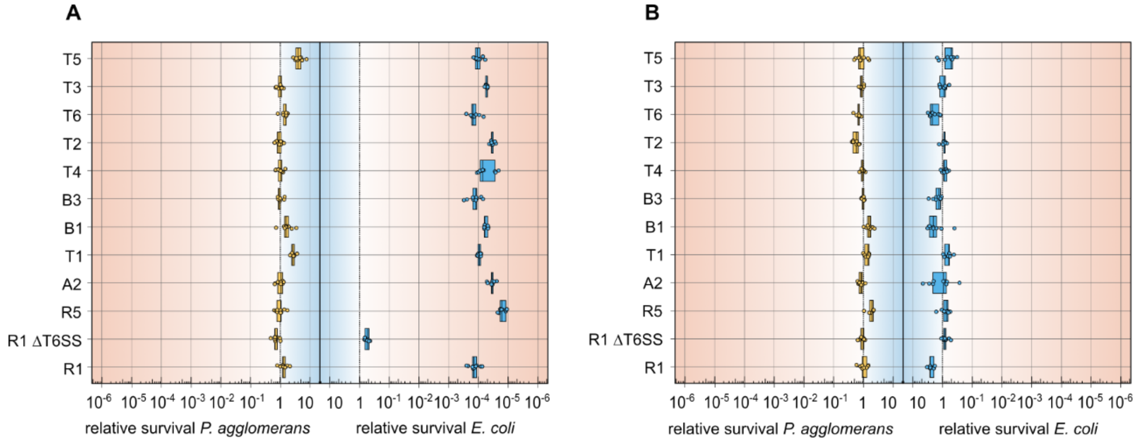


**Supplementary Figure S3**. Survival competitions of *P. agglomerans* (orange) strains against *E. coli* (blue) on solid medium (A) and in liquid medium (B). (A) All eleven *P. agglomerans* strains were equally able to severely reduce *E. coli* colony forming units (CFU) below the limit of observation when co-incubated on solid medium. Since all *Pantoea* strains reduced *E. coli* CFUs below the observable limits, *E. coli* population sizes were set to 1x10^4 cells per milliliter the respective limits of observation in our experimental set-up for the calculation of relative fitness. The T6SS Cluster 1 deficient knock-out mutant R1 ΔT6SS showed no killing against *E. coli* when co-incubated on solid medium. (B) When co-incubated in liquid medium no killing effects could be observed. This served as evidence that the killing effect of *Pantoea* against *E. coli* observed on solid medium was mediated by the Type VI secretion system of the *Pantoea* strains.

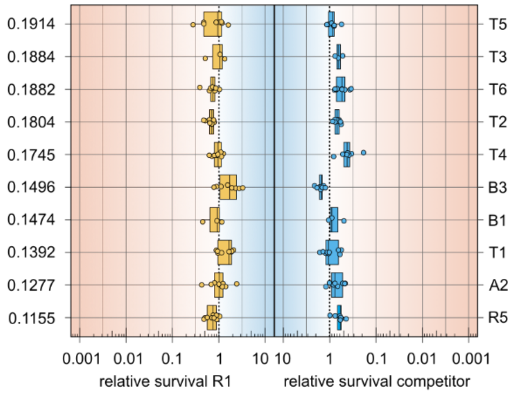


**Supplementary Figure S4**. Survival competition in liquid medium against the pan-genome jaccard distance. In liquid medium no reduction in relative survival could be observed. The fact no reduction of relative survival could be observed in liquid medium, served as evidence that reduction of relative survival observed in competitions on solid medium were caused by T6SS activity.
